# CIC-DUX4 expression drives the development of small round cell sarcoma in transgenic zebrafish: a new model revealing a role for ETV4 in CIC-mediated sarcomagenesis

**DOI:** 10.1101/517722

**Authors:** Sarah Watson, Genevieve C. Kendall, Dinesh Rakheja, Matthew E. McFaul, Bruce W. Draper, Franck Tirode, Olivier Delattre, James F Amatruda

## Abstract

CIC-DUX4 sarcoma is a rare subtype of sarcoma characterized by a devastating prognosis and resistance to conventional therapeutic strategies. So far, only few models of the disease have been reported, and its biological mechanisms remain to be elucidated. We established mosaic transgenic zebrafish expressing the human *CIC-DUX4* fusion under the control of the β-actin promoter. CIC-DUX4 transgenic fish rapidly developed aggressive soft tissue tumors with a high penetrance. RNAseq profiling revealed that fish tumors shared major common targets with human tumors and cell lines, including the overexpression of the Pea3 transcription factors, *etv4* and *etv5*. Tumor development was strongly impaired in *etv4*-deficient zebrafish, implicating Etv4 as a critical effector of CIC-DUX4-mediated oncogenesis. Altogether, we report here the first *in vivo* model of CIC-DUX4 sarcoma in zebrafish, which will represent a major tool for future preclinical research.

## Statement of significance

We report here the first successful zebrafish transgenic model of CIC-DUX4 sarcoma. Tumors arising in this model shared significant clinical and biological features with the human disease. This study highlights for the first time the oncogenic properties of CIC-DUX4 *in vivo* and the role of ETV4 in tumor development.

## Introduction

CIC-DUX4 sarcomas (CDS) consist of a group of small round cell sarcomas affecting mostly young adults and characterized by the development of aggressive soft tissue tumors with devastating prognosis and resistance to conventional therapeutic strategies (1). From a pathological point of view, CDS present with small to medium-sized, round to ovoid undifferentiated tumor cells, often organized in nodular patterns, and showing variable expression of CD99 (2). Due to their clinical and pathological partial overlap with Ewing sarcomas, CDS have long been classified as “Ewing-like sarcomas” since no evidence of FET-ETS fusion gene could be identified in those tumors. *CIC-DUX4* is a fusion of the transcriptional repressor CIC with the double homeobox protein DUX4. *CIC-DUX4* fusions were first identified in 2006, and later shown to result either from a t(4;19)(q35;q13) or a t(10;19)(q26.3;q13) translocation (3, 4), with the *DUX4* gene being located both on chromosome 4 and chromosome 10 subtelomeric regions. The resulting CIC-DUX4 fusion protein retains most of the wild-type CIC, with its DNA-binding domain fused to the C-terminal portion of DUX4, and it has thus been hypothesized that the chimeric protein would act as an aberrant transcription factor.

Additional studies have shown that CDS are biologically distinct from classical FET-ETS-positive Ewing sarcomas, and characterized by a specific transcriptomic signature (5) which includes expression of the PEA3 transcription factors ETV4, ETV5 and ETV1 (3, 6). ETV4 expression has recently been recognized as a specific diagnostic marker for the identification of CIC-rearranged sarcomas in clinical specimens (7, 8). So far, the mechanisms by which CIC-DUX4 regulates gene expression to promote oncogenesis are still poorly understood, mostly due to the rarity of the disease and the lack of appropriate *in vitro* and *in vivo* models. CIC-DUX4 was shown to be able to bind the promoters of PEA3 transcription factors and activate their expression *in vitro* (3). The oncogenic properties of the fusion protein have been more recently studied in an *ex vivo* model, showing that *CIC-DUX4* expression could transform mouse mesenchymal progenitor cells and drive the development of small round cell tumors in mice (9). However, the cell of origin of the tumor is still unknown which has hindered the development of robust animal models, and the role of ETV4 overexpression and its contribution in CIC-DUX4-mediated oncogenesis remains to be elucidated.

We report here the successful generation of the first transgenic animal model of CDS. By creating mosaic transgenic animals expressing human *CIC-DUX4* under the control of the *β-actin* ubiquitous promoter, we show that *CIC-DUX4* expression is sufficient to induce the development of small round cell sarcomas in adult zebrafish with a high penetrance. Zebrafish tumors were similar to human CDS both from pathological and biological points of view, with the expression of a common set of genes including Ets transcription factors. By using an *etv4*-deficient fish strain, we demonstrated that Etv4 is crucial to the development of CDS in zebrafish, since *etv4*-deficiency abrogated tumor development. This robust model highlights for the first time the oncogenic properties of CIC-DUX4 *in vivo* and leads the way to the design of new therapeutic strategies.

## Results

### CIC-DUX4 is a potent oncogene in transgenic zebrafish

To study the oncogenic properties of CIC-DUX4 *in vivo*, we cloned the human *CIC-DUX4* fusion together with the GFP-2A fluorescent reporter under the control of the *β-actin* ubiquitous promoter into the Tol2 transposon system (Figure 1A). The transgene was injected in one-cell-staged wild-type zebrafish embryos to be integrated in the genome in a stable mosaic manner. The embryos were raised to adulthood and monitored for tumor growth. CIC-DUX4 transgenic zebrafish showed the development of aggressive tumors, starting around 50 days post-fertilization (dpf) and affecting more than 30% of injected individuals by 90 dpf, whereas no tumor was observed in control animals injected with GFP-2A alone (Figure 1B). Tumors were mostly developing rapidly from muscle, growing either in the perimuscular tissues or as exophytic masses (Figure 1C), or in the head and brain area (Figure 1D). Pathological examination of tumors revealed two main histotypes. Most tumors developing from muscles showed the presence of proliferative small round tumor cells packed in highly cellular lesions and lacking any specific line of differentiation (Figure 1E). In contrast, tumors developing in the head presented rather with more disperse tumor cells surrounded by a frequent myxoid stroma and exhibiting neural-like differentiation (Figure 1F). The latter tumors were often large and infiltrated adjacent tissues, which made it difficult to determine whether they originated from the brain or from facial structures.

**Figure 1:**
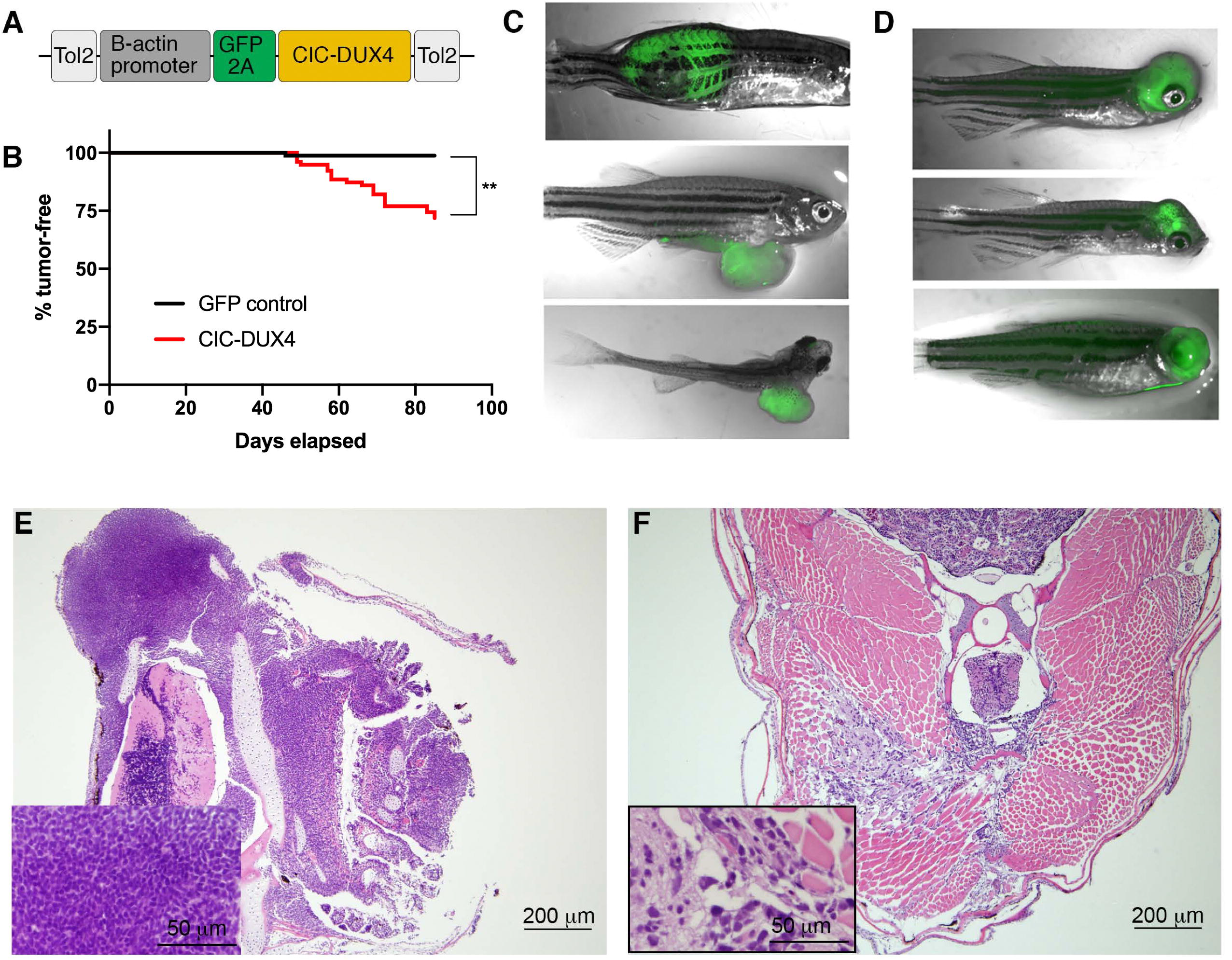
Human *CIC-DUX4* is tumorigenic in zebrafish. A: Single-cell staged wild-type zebrafish embryos were injected with mosaic GFP2A-tagged human *CIC-DUX4* under the control of the β - actin-GFP2A-CICDUX4 (CIC-DUX4) injected zebrafish (*n*=78) versus β ctin-GFP2A-pA injected controls (*n*=80) in an AB/TL wild-type genetic background. **: *p*<0.001 by Mantel-Cox Log-rank test. C and D: CIC-DUX4 induced the development of extensive tumors growing either from muscle (C) or in the head of the zebrafish (D). E and F: Representative hematoxylin and eosin stains of CIC-DUX4 induced zebrafish tumors, with either the presence of undifferentiated small round blue tumor cells (E), or tumors with a “neural-like” differentiation and myxoid stroma (F).

### CIC-DUX4 zebrafish tumors are biologically homogeneous

CIC-DUX4 may act either as an oncogene, through a gain-of-function effect via aberrant transactivation of target genes or, and non-exclusively, through a dominant negative effect on CIC’s repressor activity. To identify the genes regulated by CIC-DUX4 in zebrafish tumors, we performed whole transcriptomic analyses of 12 different CIC-DUX4 fish tumors by RNAseq. The transcriptomic signatures were compared to normal muscle from adult fish, and to other fish tumors including four malignant peripheral nerve sheath tumors (arising spontaneously in *tp53*^*M214K/M214K*^ mutant fish) and six EWSR1-FLI1-driven tumors from our previously-developed zebrafish model of Ewing sarcoma (10). Unsupervised clustering revealed that the zebrafish CIC-DUX4 form a single cluster of tumors that can be further refined in two subgoups, consistent with pathological observations of the two different histological subtypes previously observed (Figure 2A, B). Principal component analysis of the transcriptomic data similarly revealed two subgroups within the CIC-DUX4 cluster, corresponding to the histologic variants (Figure 2C).

**Figure 2:**
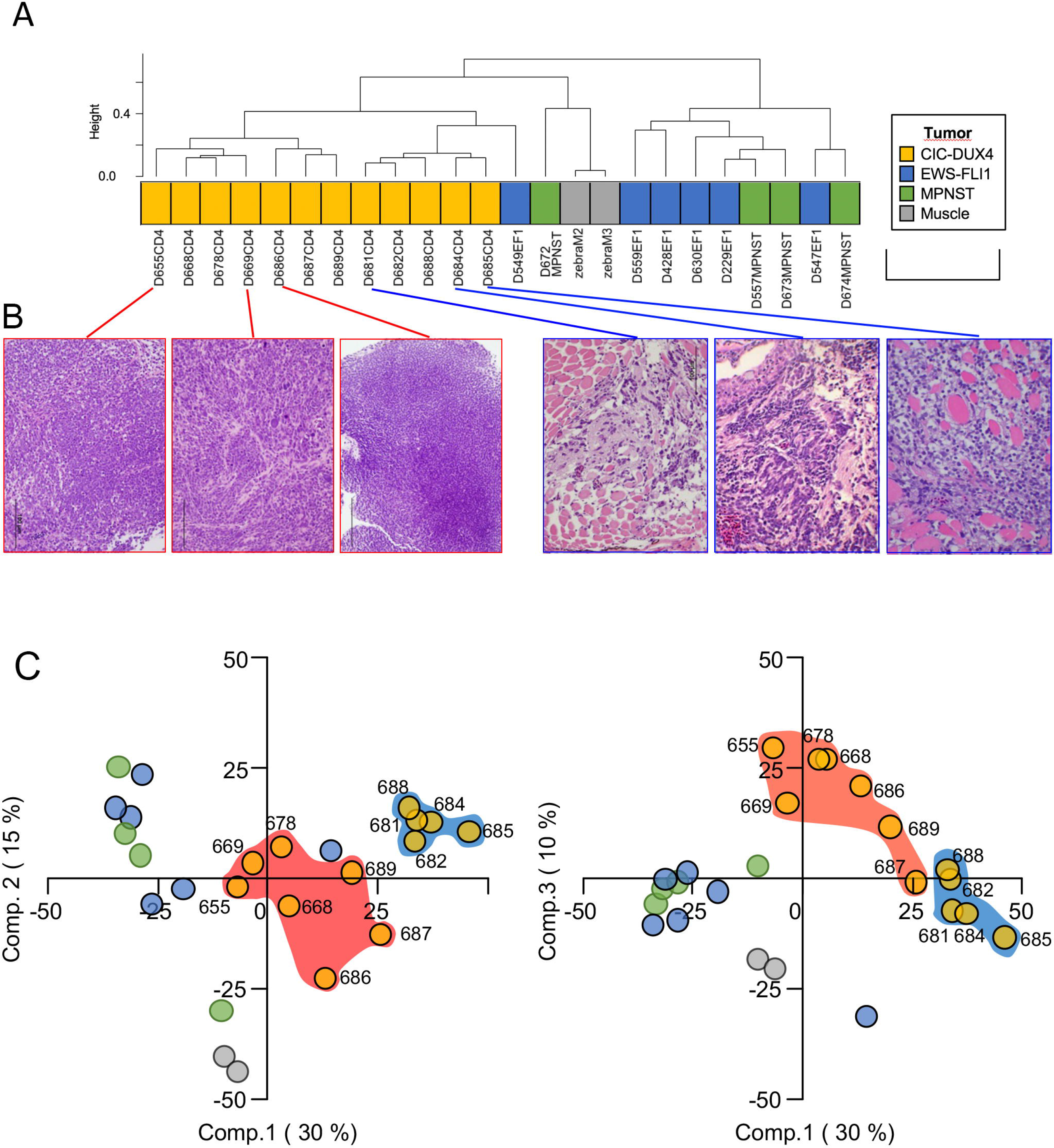
CIC-DUX4 tumors form a distinct group in unsupervised clustering analysis. Whole transcriptome analyses were performed by RNAseq on zebrafish tumors and normal tissue, including 12 CIC-DUX4 tumors (CD4), 6 EWSR1-FLI1 tumors (EF1), 4 malignant peripheral nerve sheath tumors (MPNST), and 2 samples of normal muscle from adult zebrafish.A. Unsupervised clustering showing that CIC-DUX4 tumors form a distinct group with two subgroups. B. Representative histologic images showing small round blue cell (red) and neural-like (blue) histologies. C. Principal component analyses confirmed the separation of zebrafish CDS from the EF1, MPNST or muscles samples and a further dichotomy between the 2 morphologies (highlighted by the blue and red frames, in accordance with panel B). The left panel shows the 2 first components while the right panel shows the 1st and 3rd components (component’s contribution to global variation is indicated under brackets).

### CIC-DUX4 zebrafish tumors show biological overlap with human tumors and cell lines and overexpress orthologs of the ETS transcription factors *ETV4*, *ETV5* and *ETV1*

To study the similarity of CIC-DUX4 zebrafish tumors to the human malignancies, we next compared the transcriptomic profiles of CIC-DUX4 fish tumors with those of human primary CDS. We first identified 1412 genes differentially expressed in fish CDS as compared to EF1 and MPNST tumors (Supplementary Table S2), among which 892 were up-regulated and with a defined human orthologue. Matching these 892 genes to the 595 genes specifically expressed in human CDS (6) and having a zebrafish orthologue, highlighted 97 genes commonly up-regulated in both species (Fisher exact test *P*-value < 1e^−12^).

To further refine the comparison of fish and human tumors, we compared the expression profiles of the CIC-DUX4 fish tumors with the genes regulated by CIC-DUX4 in the IB120 cell line. The IB120 cell line is derived from a primary human CIC-DUX4 tumor. CIC-DUX4 expression was inhibited *in vitro* by siRNA, and Affymetrix microarrays were performed on siCIC-DUX4 IB120 and siControl IB120 cells to identify the genes regulated by the fusion oncogene (Supplementary Table S1). Gene Set Enrichment Analyses showed a significant concordance between the genes up-regulated in the CIC-DUX4 zebrafish tumors as compared to normal muscle and the genes up-regulated by CIC-DUX4 in the IB120 cells (genes down-regulated by CIC-DUX4 inhibition) (Figure 3A). Reciprocally, genes down-regulated in fish tumors were significantly concordant with the genes down-regulated by CIC-DUX4 (up-regulated upon CIC-DUX4 inhibition) (Figure 3B), showing that the fusion oncogene similarly regulates common targets in fish tumors and in the human primary tumor-derived cell line.

Comparing the 97 genes commonly up-regulated in fish and human tumors with the 636 genes upregulated by CIC-DUX4 in the IB120 cell line (Supplementary Table S1) and having a known zebrafish orthologue, we identified 26 genes (exact Fisher-test *P*-value: 4e^−9^) that were overexpressed in all three models (Table 1), which included the PEA3 transcription factors ETV4 and ETV5.

**Figure 3:**
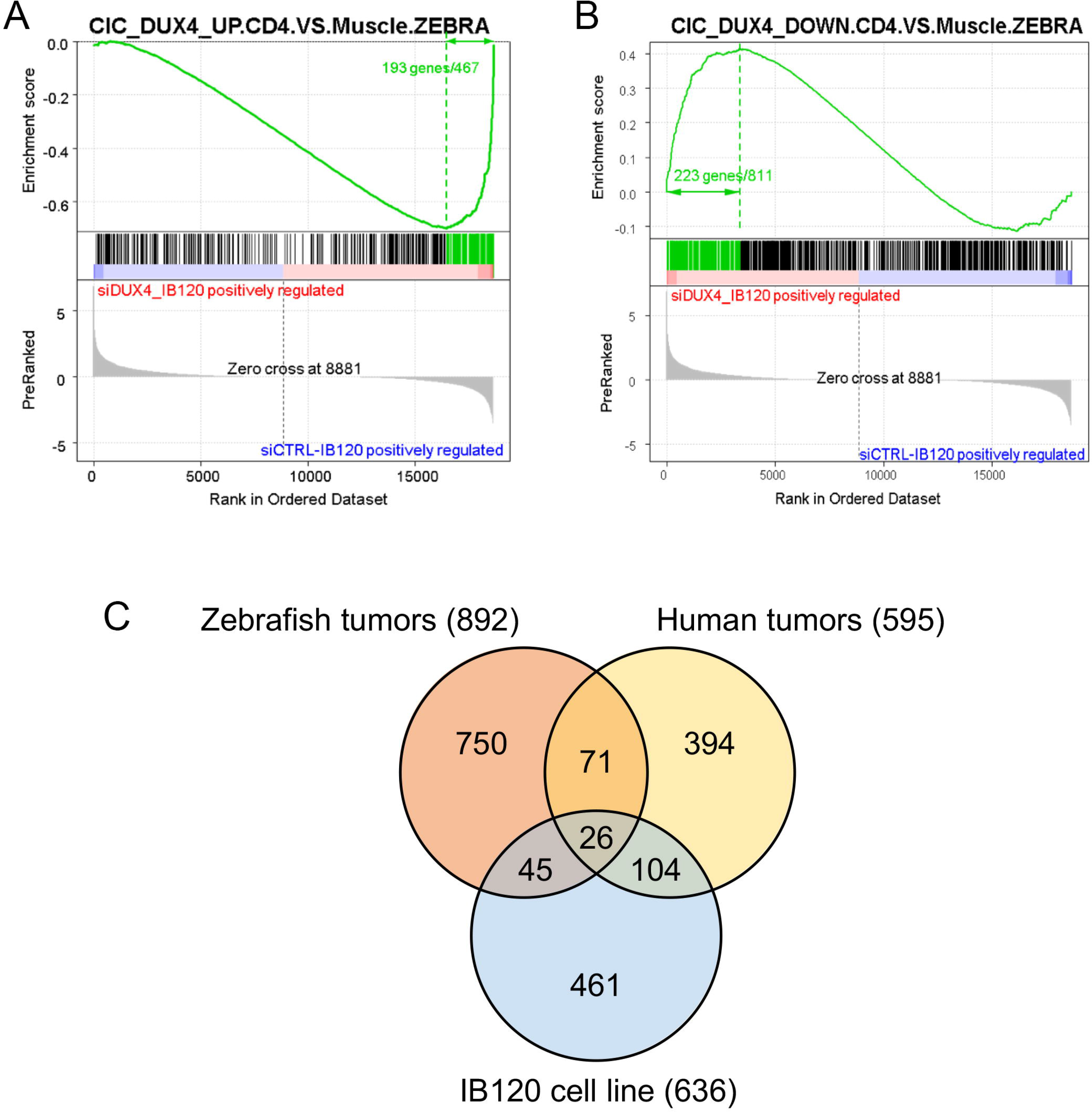
Cross-species oncogenomics in zebrafish tumors, human tumors, and cell line. A: Gene Set Enrichment Analyses show a significant concordance between genes up-regulated in CIC-DUX4 tumors versus normal tissue and genes up-regulated by CIC-DUX4 in the IB120 cell line (siCTRL_IB120 positively regulated). B: Gene Set Enrichment Analyses show a significant concordance between genes down-regulated in CIC-DUX4 tumors versus normal tissue and genes down-regulated by CIC-DUX4 (siDUX4_IB120 positively regulated) C: Intersection of genes up-regulated in human and fish tumors and by CIC-DUX4 in the IB120 cell line identified 26 genes in common to the three models, including the PEA3 transcription factors ETV4 and ETV5.

**Table 1:**
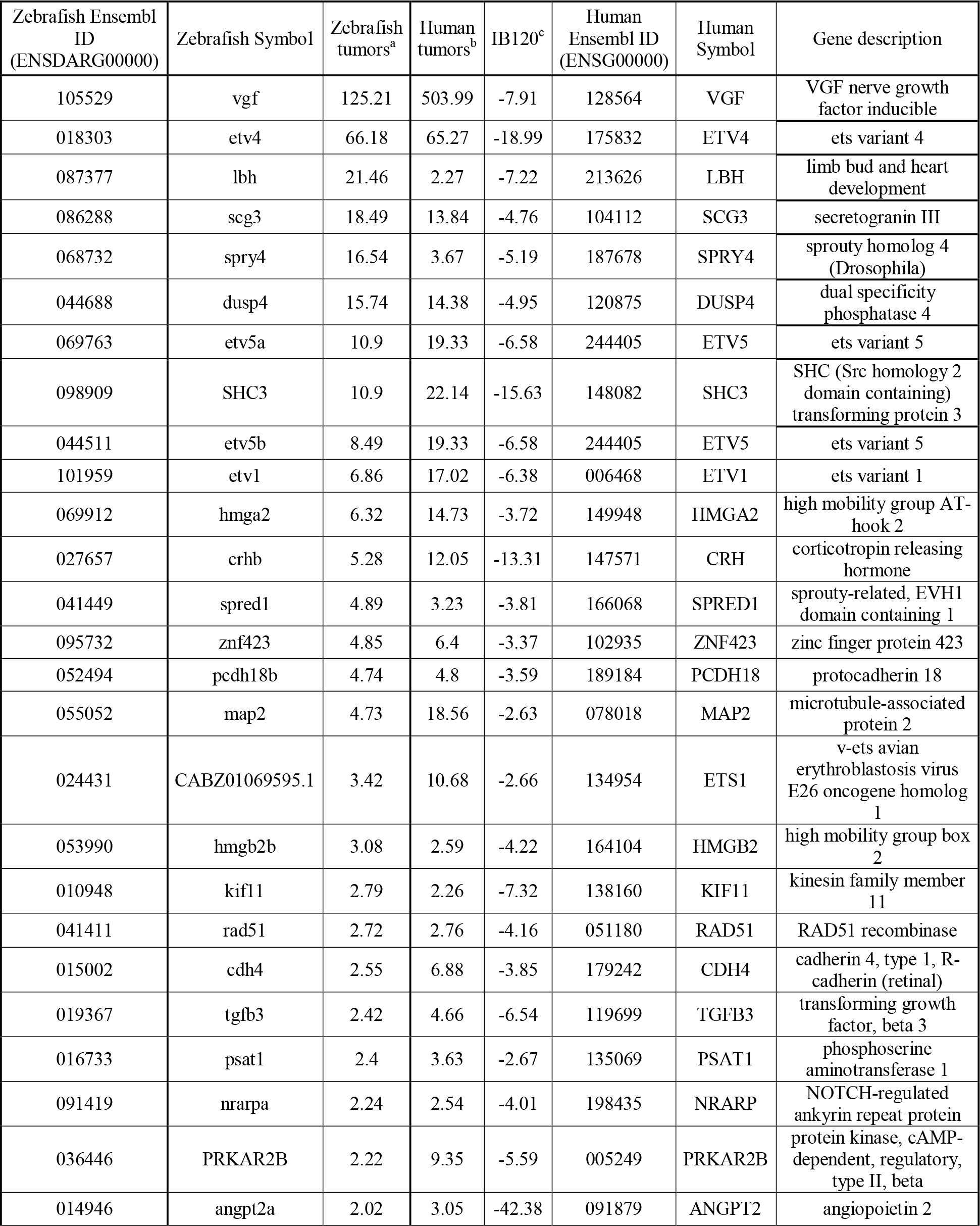
The 26 genes similarly regulated in human and fish tumors and modulated by CIC-DUX4 in the IB120 cell line

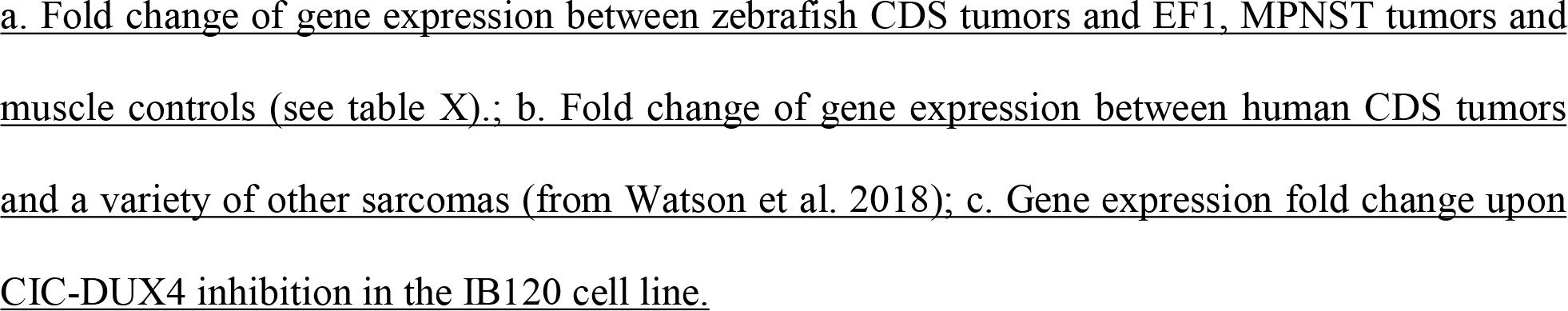

### Etv4 expression is necessary for CIC-DUX4 mediated oncogenesis in vivo

ETV4 overexpression has already been well-documented in CDS; however its role in CIC-mediated sarcomagenesis has never been studied *in vivo* so far. To address this question, we took benefit of an *etv4*-deficient fish strain, in which *etv4* has been knocked-out by CRISPR-Cas9 technology (*etv4-/-).* Human CIC-DUX4 was integrated in the genome of wildtype or *etv4-/-* fish using the Tol2 transposon system to generate mosaic transgenic animals, and adult fish were monitored for tumor development. As previously shown, CIC-DUX4-driven tumor development was highly penetrant in the wild-type background with more than 30% of fish developing tumors by 75 dpf. In contrast, *etv4* deficiency significantly impaired tumor formation with less than 10% of *etv4-/-* fish developing tumors in the same time-frame (Mantel-Cox Log-rank test *P*-value: 0.02) (Figure 4), showing that Etv4 is major mediator of CIC-DUX4-induced sarcomagenesis *in vivo*.

**Figure 4:**
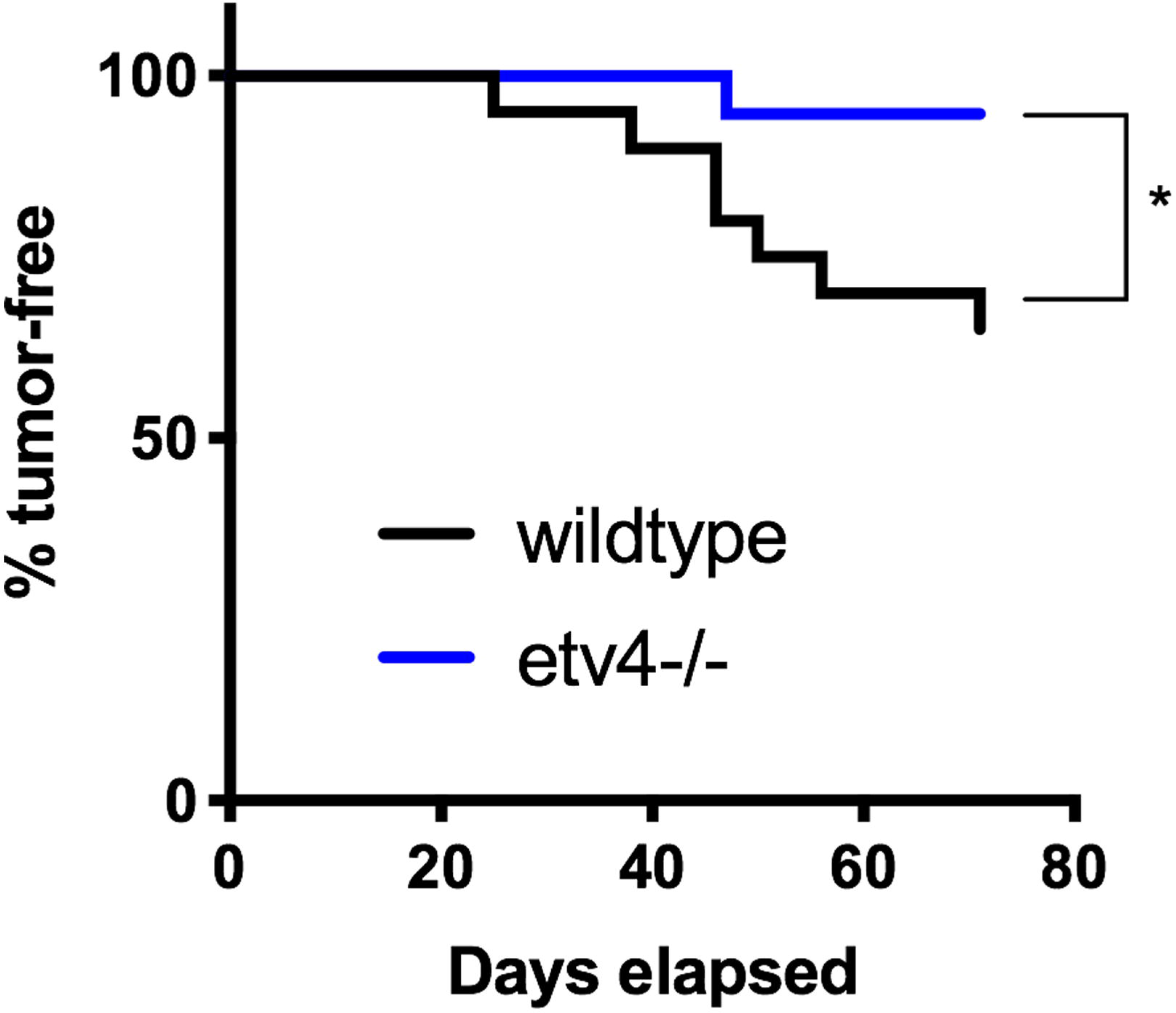
ETV4 is required for CIC-DUX4-mediated oncogenesis. β-actin-GFP2A-CICDUX4 was injected into AB/TL wildtype (*n*=19) and ETV4-deficient (etv4-/-; *n*=18) zebrafish embryos, and adult fish were monitored for tumor development. ETV4-deficiency significantly suppressed tumor growth compared to AB/TL wild-type controls. *: *p*=0.02 by Mantel-Cox Log-rank test.

## Discussion

Since the first identification of CIC-DUX4 fusion transcript in two human small round cell sarcomas in 2006, more than 150 different cases have been reported in the literature (1, 4, 11). The advances in the techniques of molecular biology and the identification of specific markers to aid in pathological diagnosis have led to a significant increase in the number of reported cases (8, 12–14), and it is now acknowledged that CDS represents the most frequent subtype of the previously-called “Ewing-like” tumors. Despite evidence that CDS are biologically distinct from classical Ewing sarcoma and respond poorly to conventional chemotherapy, most patients are still treated in the same way and included in the same clinical trials as Ewing sarcoma patients.

So far, the functions of *CIC-DUX4* fusion gene have been mostly studied *in vitro*. *CIC* encodes an HMG-box transcriptional repressor homologous to Drosophila *capicua* (15). Although several breakpoints within *CIC* and *DUX4* have been reported in CDS (16), the resulting chimeric proteins always keep the majority of CIC, including its DNA-binding domain, fused together with the C-terminal part of DUX4. In the original study describing the first cases of CDS, *in vitro* experiments showed that CIC-DUX4 could transform NIH3T3 cells (3). Moreover, in U2OS osteosarcoma cells, the chimeric protein acted as a strong transcriptional activator, with the binding of the promoter and the resulting up-regulation of most of its target genes (including ETS transcription factors ETV5 and ETV1). Other *in vitro* models have been reported, including the generation of cell lines generated from patients-derived xenografts (17). More recently, Yoshimoto and colleagues reported an *ex-vivo* model of CDS, with the successful development of small round cell sarcoma in mice xenografted with mouse embryonic mesenchymal cells previously transduced by CIC-DUX4 (9). In this model, tumor latency was of less than a month, the penetrance was 100% and systemic diffusion of the disease frequent, reminiscent of the aggressiveness of human disease.

Our model is the first transgenic animal model of CDS. We demonstrated that CIC-DUX4 expression is sufficient to drive the development of tumors in transgenic zebrafish with a remarkable penetrance for a mosaic model. Moreover, the use of a ubiquitous promoter was chosen due to the fact that CDS cell of origin is unknown; thus the expression of the oncogene under the control of an appropriate tissue specific promoter might even increase tumor penetrance. Transgenic zebrafish tumors arose mainly from muscle or from the head with a very short latency and developed as undifferentiated aggressive malignancies with invasion of local anatomical structures. This clinical and pathological behavior is similar to human CDS, with most tumors developing from limbs or abdominal wall muscles and growing extensively. In our model, tumors arising from the head or the brain presented a rather different pathological aspect, with increased neural differentiation and myxoid stroma. Whether the pathological differences observed within fish tumors reflect the cell of origin of the disease remains to be further studied. Of note, head and neck CDS are not uncommon in humans (18), and several cases of CDS arising from the brain have been reported, sometimes with atypical histology (19, 20). Moreover, the related CIC-NUTM1 fusion is a recurrent genetic lesion in a subset of central nervous system Primitive Neuroectodermal Tumors (CNS-PNETs) (21, 22).

The overexpression of Ets transcription factors has long been identified as a hallmark of CDS, and ETV4 immunostaining is even used in clinical practice to help diagnosis (7). PEA3 transcription factors ETV1, ETV4 and ETV5 are expressed in numerous organs during embryonic development and in adults, and are thought to play various physiological functions involved in morphogenesis of limbs and kidneys, and in motor coordination (23). In oncogenesis, EWSR1-ETV1 and EWSR1-ETV4 fusions have been reported as rare events in classical Ewing sarcoma (24), and *TMPRSS2* fusions with either *ETV1*, *ETV4* or *ETV5* are also observed in prostate carcinoma (25). Moreover, *ETV4* overexpression has been identified as an independent factor of poor prognosis in various types of malignancies, including breast, gastric, colorectal, and lung carcinoma (23). Under physiological conditions, CIC transcriptionally represses the expression of *ETV4* and downstream targets, including genes involved in extracellular matrix remodeling and cell proliferation (26). Importantly, oligodendroglioma harboring *CIC* inactivating mutations present *ETV4* overexpression and are characterized by an aggressive behavior (27). And inactivation of *CIC* in lung and gastric carcinoma result in *ETV4* overexpression and increased risk of metastasis (28). Thus, whether *ETV4* overexpression in CDS is the direct consequence of CIC-DUX4 transcriptional activation or might result from the loss of function of wild-type *CIC* remains to be further studied *in vivo*. We showed however that Etv4 is crucial to tumor development in our model, since CIC-DUX4-mediated tumor growth was abrogated in Etv4-deficient zebrafish. This finding is of major interest for the clinical management of CDS, for which conventional therapies are poorly effective, especially as strategies to pharmacologically target Ets transcription factors are being developed. We thus believe that this first *in vivo* model of CDS will be of particular interest to further study the biological mechanisms of the disease and will constitute a major tool for the elaboration of new targeted therapies for the human disease.

## Methods

### Zebrafish husbandry

*Danio rerio* were maintained in an aquatics facility according to industry standards. Vertebrate animal work was carried out according to protocols approved by the Institutional Animal Care and Use Committee at UT Southwestern, an AALAC-accredited institution committee. Zebrafish were originally obtained from the Zebrafish International Resource Center (https://zebrafish.org). The wildtype lines used were AB/TL hybrids. The *tp53* mutant line, *tp53*^*zdf1*^, was a kind gift from Tom Look (29). The *etv4* mutant line was generated in the wildtype NHGRI-1 strain.

### Generation of a Tol2 enabled CIC-DUX4 expression construct

The human *CIC-DUX4* coding sequence was sub-cloned directly from a patient tumor and integrated into the Gateway expression system (Thermo) by adding 5’ and 3’ ATT sites (attb2r/attb3) with primers and high-fidelity PCR. Purified PCR products were cloned into a 3’ entry clone as previously described (30). The tol2 kit β-actin promoter, 3’ SV40 late poly A signal construct, and pDestTol2pA2 destination vector were used for construct generation and expression in zebrafish (31). The GFP viral 2A sequence was a kind gift from Steven Leach (32), and was previously sub-cloned into a middle entry Gateway expression system (33). Tol2 mRNA was synthesized from pCS2FA-transposase from Koichi Kawakami (34). The constructs utilized in this study include: β-actin-GFP2A-*CICDUX4*.

### CIC-DUX4 integration into zebrafish embryos and characterization of tumor incidence

Zebrafish were injected at the single cell stage with injection mixes containing 50ng/μL Tol2 transposase mRNA, 40ng/μL of BetaActin-GFP2A-pA or BetaActin-GFP2A-*CICDUX4* Tol2 enabled plasmid, 0.1% phenol red, and 0.3X Danieau’s buffer. The same microinjection needle was utilized to inject wildtype and etv4-/-mutant zebrafish to compare *CIC-DUX4* tumorigenic capacity. Zebrafish presenting with tumors were screened under the fluorescent microscope to determine if they were GFP positive, were collected, and the presence of malignancies confirmed by H&E staining and visual review by a pathologist.

### Zebrafish tumor collection and downstream processing of RNA and histology

Zebrafish with tumors were euthanized and screened under a Nikon SMZ25 fluorescent stereomicroscope to detect GFP expression indicative of CIC-DUX4 expression. Fresh GFP+ tumor tissue was resected and snap frozen in liquid nitrogen. Total RNA was isolated from zebrafish using the TRIzol reagent kit (Invitrogen) and was utilized for RNAseq. The remaining tumor specimen was fixed in 4% paraformaldehyde/1XPBS for 48 hours at 4°C (Fisher). Samples were then de-calcified in 0.5M EDTA for 3-5 days and mounted in paraffin blocks for microtome sectioning. Hematoxylin and eosin (H&E) staining was performed on de-paraffinized slides.

### Generation of etv4 mutant allele by CRISPR-Cas9

A single-stranded guide RNA and Cas9 mRNA were prepared as described (35). The guide RNA was designed to target the second of exon of etv4 (5’-<u>aattaatacgactcactata</u>GGACGCAGAATTACCCCCTC*gttttagagctagaaatagc*-3’; T7 promoter underlined, *etv4* target sequence uppercase, overlap to sgRNA scaffold oligo italicized). An injection solution was prepared with 12ng/μL of guide RNA, 30ng/μL Cas9 mRNA, 300mM KCl, and 0.1% phenol red. 2nL of injection solution was injected into the yolk of 1-cell stage zebrafish embryos. Cutting efficiency of the guide RNA was determined by High Resolution Melt Analysis (HRMA) at 24hpf with 8 injected embryos and 8 uninjected controls using the primers etv4-Fwd (5’-AGAGGTCGCAAGGAAATGGGC-3’) and etv4-Rev (5’-AACTTGCTTGTTATTTTACCTTCAGAT-3’)(36). Sperm from injected males was screened by HRMA for germline transmission of etv4 alleles with indels. A 14bp frameshift-causing deletion was identified in an injected male founder by Sanger sequencing of genomic DNA. The frameshift allele was confirmed by sequencing an etv4 cDNA from outcrossed embryos.

### Cell line and reagents

IB120 cells are a kind gift from Dr. Jean-Michel Coindre and Frederic Chibon (Institut Bergonié, Bordeaux, France) and were derived from a primary tumor harboring the *CIC-DUX4* fusion. IB120 cells were cultured in Roswell Park Memorial Institute (RPMI)-1640 medium (Life Technologies, Saint Aubin) supplemented with 10% fetal calf serum (Eurobio, Courtaboeuf) and 50U/50μg penicillin/streptomycin (Life Technologies). FlexiTube siRNAs against *DUX4* and control siRNA were purchased from Qiagen (Courtaboeuf). Transfections of siRNAs were performed using the Lipofectamine RNAiMAX transfection reagent (Life Technologies). 150 000 cells were seeded in 6-well plated in 2 mL of antibiotics-free medium and transfected 24 hours later with 20nM of total siRNA. RNA extraction was performed 48 hours after transfection with RNA extraction kits (Macherey) following manufacturer’s instructions and then used for Affymetrix microarrays.

### Microarray

HG-U133-Plus2 GeneChip microarrays were performed by the Affymetrix platform of the Institut Curie according to the Affymetrix GeneChip Expression Analysis Technical Manual. Expression profiles were normalized using the gcrma package version 2.34.0 in the R 3.0.2 environment (R Development Core Team, 2012) with Brainarray Entrez gene CDF v18. Quality assessment was based on relative log expression and normalized unscaled standard errors. Raw data were deposited on the Gene expression Omnibus database, as part of the <u>GSE60740</u> dataset:(https://www.ncbi.nlm.nih.gov/geo/query/acc.cgi?acc=GSE60740).

### Paired-end RNA sequencing

Library constructions were performed following the TruSeq Stranded mRNA LS protocol (Illumina, San Diego, CA, USA) and sequencing was performed on HiSeq 2500 (50nt paired-end) Illumina sequencing machines, with an average of 30 million of reads per sample, by the NGS platform of the Institut Curie. FASTQ files will be deposited on the Gene expression Omnibus database.

### Bioinformatic Analysis

Gene expression values were extracted by Kallisto v0.42.5 [Bray NL, Pimentel H, Melsted P, *et al.* Near-optimal probabilistic RNA-seq quantification. *Nat Biotechnol* 2016; **34**: 525-527] with GRCz10 release 90 genome annotation. Clustering and PCA were computed using the R packages Cluster v2.0.3 and FactoMiner v1.31.4, respectively. Gene ontology analyses were performed online (https://david.ncifcrf.gov/) with DAVID v6.7 tool [Huang DW, Sherman BT, Lempicki RA. Bioinformatics enrichment tools: paths toward the comprehensive functional analysis of large gene lists. *Nucleic Acids Res* 2009; **37**: 1–13]. Differential expression analysis were performed using Welsh t-tests and the associated P-values were corrected by the Bonferroni procedure. Only the genes with a Bonferroni corrected P-value under 0.05 and an absolute fold change above 2 were further considered. Orthologues were identified using a combination of annotations from Ensembl (www.ensembl.org) and the Zebrafish Information Network (https://zfin.org)databases.

## Acknowledgements

Supported by grants from the Cancer Prevention and Research Institute of Texas (RP120685), the 1Million4Anna Foundation and the Curing Kids Cancer Foundation to JFA; the National Institute of Child Health and Development (R01 HD081551) to BWD; and the National Institute of General Medical Science (T32 GM007377) to MEM. This work was also supported by grants from the Institut Curie, the Inserm, the European PROVABES (ERA-649 NET TRANSCAN JTC-2011), ASSET (FP7-HEALTH-2010-259348) projects and the following associations: Courir pour Mathieu, Dans les pas du Géant, Les Bagouzamanon, Enfants et Santé, M la vie avec Lisa, Lulu et les petites bouilles de lune, les Amis de Claire, l’Etoile de Martin and the Société Française de lutte contre les Cancers et les leucémies de l’Enfant et de l’adolescent. GCK was supported by a QuadW-AACR Postdoctoral Fellowship for Clinical/Translational Sarcoma Research, a Young Investigator Grant from Alex’s Lemonade Stand, and a Hartwell Foundation Postdoctoral Fellowship. High-throughput sequencing has been performed by the NGS platform of Institut Curie, supported by the grants ANR-10-EQPX-03 and ANR10-INBS-09-08 from the Agence Nationale de la Recherche (investissements d’avenir) and by the Canceropôle Ile-de-France.

